# Immature HIV-1 Lattice Assembly Dynamics are Regulated by Scaffolding from Nucleic Acid and the Plasma Membrane

**DOI:** 10.1101/163295

**Authors:** Alexander J. Pak, John M. A. Grime, Prabuddha Sengupta, Antony K. Chen, Aleksander E. P. Durumeric, Anand Srivastava, Mark Yeager, John A. G. Briggs, Jennifer Lippincott-Schwartz, Gregory A. Voth

## Abstract

The packaging and budding of Gag polyprotein and viral ribonucleic acid (RNA) is a critical step in the human immunodeficiency virus-1 (HIV-1) lifecycle. High-resolution structures of the Gag polyprotein have revealed that the capsid (CA) and spacer peptide 1 (SP1) domains contain important interfaces for Gag self-assembly. However, the molecular details of the multimerization process, especially in the presence of RNA and the cell membrane, have remained unclear. In this work, we investigate the mechanisms that work in concert between the polyproteins, RNA, and membrane to promote immature lattice growth. We develop a coarse-grained (CG) computational model that is derived from sub-nanometer resolution structural data. Our simulations recapitulate contiguous and hexameric lattice assembly driven only by weak anisotropic attractions at the helical CA-SP1 junction. Importantly, analysis from CG and single-particle tracking photoactivated localization (spt-PALM) trajectories indicates that viral RNA and the membrane are critical constituents that actively promote Gag multimerization through scaffolding, while over-expression of short competitor RNA can suppress assembly. We also find that the CA amino-terminal domain imparts intrinsic curvature to the Gag lattice. As a consequence, immature lattice growth appears to be coupled to the dynamics of spontaneous membrane deformation. Our findings elucidate a simple network of interactions that regulate the early stages of HIV-1 assembly and budding.

**SIGNIFICANCE STATEMENT:** In order for human immunodeficiency virus to proliferate, viral proteins and genomic dimers are assembled at host cell membranes and released as immature virions. Disrupting this key intermediate step in viral replication is a potential target for treatment. However, a detailed molecular view of this process remains lacking. Here, we elucidate a network of constitutive interactions that regulate viral assembly dynamics through a combined computational and experimental approach. Specifically, our analysis reveals the active roles of nucleic acid and the membrane as scaffolds that promote the multimerization of Gag polyprotein which proceeds through multi-step and self-correcting nucleation. Our findings also illustrate the functional importance of the N-terminal, C-terminal, and spacer peptide 1 protein domains.

## INTRODUCTION

Human immunodeficiency virus type 1 (HIV-1) spreads infection through a robust replication cycle. A critical step during this lifecycle is the assembly and packaging of viral proteins and ribonucleic acid (RNA) at the plasma membrane interface, followed by the budding and release of immature virions (1-4). Understanding the molecular mechanisms that regulate this process may highlight new opportunities for anti-retroviral treatment strategies.

The main structural constituent throughout HIV-1 replication is the Gag (group specific antigen) polyprotein (5, 6), which comprises around 50% of the viral particle mass (7). Gag contains several functional domains: the matrix (MA) domain for membrane association, the capsid (CA) domain for protein-protein association, the nucleocapsid (NC) domain for RNA association, the p6 domain for ESCRT machinery recruitment, and the spacer peptide 1 (SP1) and 2 (SP2) domains. Cryo-electron tomography (cryoET) experiments have demonstrated that Gag polyproteins form incomplete spherical and hexameric lattices within immature virions and arrested budding sites (7-11). The large gap-like defects throughout the immature lattice are hypothesized to accommodate the strain from lattice curvature (10), in direct contrast to the incorporation of pentameric units that permit curvature in the closed mature capsid core (12-17).

Resolving the structure of multimerized Gag constructs within immature virions and virus-like particles (VLPs) has provided many insights into the relationship between polyprotein structure and supramolecular assembly. Most notably, recent advances in cryo-electron microscopy (cryoEM) and cryoET have revealed an unprecedented sub-nanometer resolution of Gag-Gag interfaces (18-21). From these studies, important interactions have been inferred from close protein-protein contacts, such as at the CA^CTD^ helix 9 interface for dimerization and the helical CA-SP1 junction interface for hexameric bundling. Biochemical experiments have corroborated these findings; for example, the importance of the SP1 domain and nearby motifs, including the major homology region and type II β-turn, was demonstrated by the production of aberrant virions following mutations to associated residues (21-23). In fact, the helical transition of the CA-SP1 junction has been proposed as a conformational molecular switch that regulates immature lattice assembly (21-24).

For efficient virion production, concerted interactions between Gag and both RNA and lipids in the cell membrane appear to be necessary. *In vitro* assembly of VLPs with immature-like spherical geometries requires, at a minimum, Gag binding to oligonucleotides (25, 26) or through leucine zipper motifs (27) while binding to inositol derivatives appears to adjust particle sizes (28). Recent studies have suggested that Gag assembly at the cell membrane can promote selective RNA dimerization and packaging (29, 30); competitor tRNA is also observed to positively-regulate viral RNA selectivity (30). In contrast, over-expression of non-silencing microRNA is suggested to negatively-regulate viral production by impeding Gag assembly (31). The composition of RNA within virions has also been connected to particle size dispersity (32). Furthermore, lipid domains in the plasma membrane may mediate Gag targeting and assembly (33, 34) while membrane curvature can initiate nucleation of ESCRT machinery for scission (35). It is therefore clear that a network of interactive processes between Gag, RNA, and the plasma membrane may coordinate viral replication. Nonetheless, the underlying mechanisms that dictate the interfacial assembly dynamics remain unclear.

Molecular dynamics (MD) simulations may afford new insight into the dynamics of these cellular processes. As opposed to very computationally demanding all-atom MD simulations, coarse-grained (CG) MD models are particularly attractive as a means to access biological phenomena that typically occur on very large length and time-scales. For example, CG simulations have recently predicted fundamental mechanisms in mature CA assembly (15, 17) and MA domain anchoring to the lipid membrane (36).

Emerging fluorescence techniques, such as total internal reflection fluorescence (TIRF) and single-particle tracking photoactivated localization microscopy (spt-PALM), also enable the probing of protein mobility during live-cell imaging with high specificity, especially in crowded and heterogeneous environments such as viral buds (37, 38). Recently, spt-PALM analysis revealed a notable suppression in mobility for Gag clustered within viral puncta compared to membrane-bound or cytosolic Gag (31, 37). Hence, cooperative understanding from computer simulations and fluorescence experiments may help elucidate assembly dynamics at the cell membrane interface.

In this work, we present a CG computational study with our modeling developed with the help of recent experimental structures. We also present spt-PALM experimental data of Gag self-assembly during the early stages of immature lattice growth. Our intention here is to shed critical light on the network of constituents and interactions required for viable self-assembly and budding. First, we use CG MD simulations to investigate the assembly of ΔCA^NTD^-CA^CTD^-SP1 Gag and find that nucleation proceeds through six-helical bundling at the CA^CTD^-SP1 junction contingent on scaffolding from RNA and the membrane. In particular, we focus on the role of RNA through analysis of Gag diffusion and clustering based on spt-PALM and CG MD trajectories. Interestingly, these lattices, while contiguous and hexameric, are predicted to be flat and hence, unlikely to instigate budding. Next, we introduce the CA^NTD^ domain and find that the Gag lattice exhibits an intrinsic curvature. Simulations of these Gag constructs reveal that spontaneous curvature of the membrane promotes efficient nucleation, while gradual membrane deformation is essential for extended CA-SP1 lattice growth. Taken together, these findings provide insight into molecular-scale mechanisms that regulate the packaging and budding of immature HIV-1 virions.

## RESULTS

As depicted in Figure 1(a), we have developed a CG model of the CA-SP1 dimer that is directly derived from experimental structural data of the immature CA^NTD^-CA^CTD^ (Protein Data Bank (PDB) Accession 4USN (19)) and CA^CTD^-SP1 (PDB 5I4T (21)) domains. The CG model is designed to recapitulate the internal structure of the polyprotein, relevant protein-protein interfaces, and quaternary structures (Figure 1(b)) through a protocol adapted from previous work on the mature CA (17). Importantly, the collective α-helices within each of the three protein domains are maintained as rigid bodies (i.e., six total per dimer) with excluded volume interactions, while inter-domain flexibility is maintained with a soft elastic network model. The dimerization is preserved through weak harmonic bonds at the helix 9-helix 9 (H9-H9) interface in the CA^CTD^ domain. Finally, the only attractive Gag-Gag interaction occurs between SP1 domains through virtual particles that serve as binding sites (see Figure 1(a)). In addition, we model RNA and the cell membrane as a linear chain and hexagonal mesh of CG particles, respectively. Attractive interactions are included between the top-most bead of the CA^NTD^ domain and the membrane and the bottom-most bead of the SP1 domain and RNA in order to mimic MA-membrane and NC-RNA interactions, respectively. Full details of the model are described in the Methods section.

**Figure 1.**
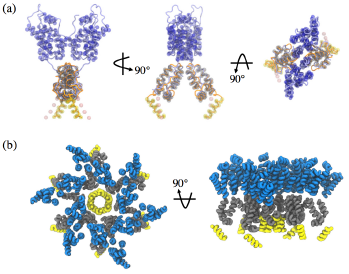
Coarse-grained representation of CA-SP1 polyprotein. (a) Schematic of all CG sites of the CA-SP1 dimer shown as beads and reference experimental structures (Refs. (19) [blue] and (21) [orange]) shown as tubes. The CA^NTD^, CA^CTD^, and SP1 particles are depicted as transparent blue, gray, and yellow spheres, respectively. The pink spheres depict virtual particles, which act as tethering sites for adjacent SP1 helices. (b) Schematic of a hexameric bundle formed from six Gag dimers. Each helix is shown as a tube with light blue, gray, and yellow denoting CA^NTD^, CA^CTD^, and SP1 domains, respectively; this convention is used for all subsequent molecular snapshots.

### Immature Lattice Assembly of ΔCA^NTD^-CA^CTD^-SP1 Gag Constructs

#### Assembly through SP1 Helical Bundling is Activated by RNA and Membrane Scaffolds

We first investigate the factors that enable self-assembly of CA^CTD^-SP1 into an immature lattice at the plasma membrane interface. Our intention is to ascertain the assembly competence of CA^CTD^-SP1 as facilitated by interactions at the CA-SP1 junction and the presence of both RNA and the cell membrane.

We prepared simulations with 144 CA-SP1 dimers initially arranged in an evenly-spaced 12×12 grid (8.5 nm separation) along the *xy* plane; final *xy* lengths fluctuated around 102 nm. The membrane and a 3500 nucleotide (nt) RNA were respectively placed 10 nm above and 20 nm below the layer of Gag dimers; here, the length of the RNA chain is deliberately long enough to provide an excess of binding sites. We also maintained membrane-CA^NTD^ interactions, which were included to implicitly represent Gag binding to the lipid bilayer, e.g., through exposed myristyl and the MA domain; the excluded volume interactions of the CA^NTD^ domain were removed to emulate its deletion. Initially, a repulsive barrier was placed between the RNA and CA-SP1 dimers to independently evolve disordered configurations of the RNA and membrane-bound Gag over 5×10^5^ CG MD timesteps (τ) at 310 K. The barrier was subsequently removed and simulations were performed for 7.5×10^7^ τ with trajectories saved every 2.5×104 τ for analysis. We note that the minimum SP1-SP1 binding strength (*E*_SP1_ = 2.35 kcal/mol) required to initiate Gag association was used for these simulations.

As seen in Figure 2(a), membrane-bound CA^CTD^-SP1 dimers readily self-assemble into a contiguous hexagonal lattice (e.g., the orientation distribution shown in the bottom inset of Figure 2(a) exhibits six-fold symmetry) once tethered to RNA. However, in the absence of binding to either membrane or RNA (e.g., top inset of Figure 2(a)), Gag multimerization was not observed throughout the entirety of our simulation trajectory. This suggests that the presence of both RNA and membrane scaffolds may be necessary for efficient immature lattice assembly. As an additional test, we subsequently removed both the RNA and membrane and found that the extended lattice persisted indefinitely. The stability of the lattice suggests that the CA-SP1 junction interactions are sufficient for multimerization while RNA and cell membranes serve as catalysts for immature lattice assembly; although *in vitro* self-assembly of Gag in solution is possible, additional agents such as nucleic acid or inositol derivatives are necessary to efficiently grow VLPs with spherical morphologies (25, 26, 28).

**Figure 2.**
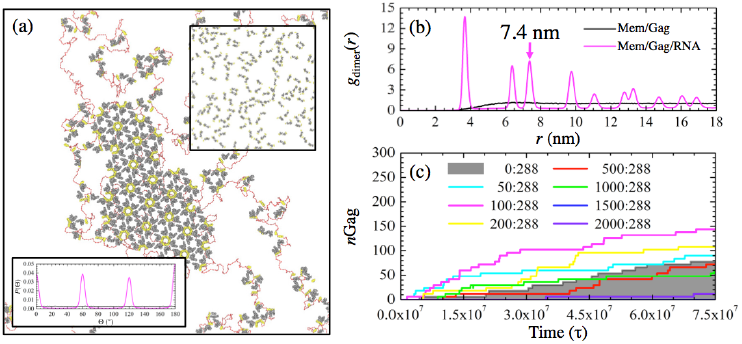
Nucleic acid modulates self-assembly of CA^CTD^-SP1. (a) Depiction of the flat immature CA^CTD^-SP1 lattice (gray and yellow tubes) at the membrane interface (not shown) with bound RNA (red chain). The top inset shows the final configuration of CA^CTD^-SP1 in the absence of RNA (and removal of the membrane yields comparably disperse Gag). The bottom inset shows the probability distribution of CA^CTD^ orientation relative to dimers within a radial distance of 8.5 nm while in the presence of RNA (purple; exhibiting six-fold symmetry) and without RNA (black). (b) Two-dimensional pair distribution function between the center of mass of Gag dimers (*g*_dimer_) as a function of radial distance (*r*). The arrow indicates the calculated lattice spacing of the hexagonal lattice. (c) Evolution in the number of Gag monomers (*n*_Gag_) within completed hexamers as a function of MD timesteps (τ). The legend lists the ratio of 20-nt miRNA to Gag monomers (i.e., [miRNA]:[Gag]) within the simulation domain. Trajectories within the shaded region exhibit hindered assembly relative to the miRNA-absent case.

Remarkably, the assembled lattice derived from our simple CG model is predicted to preserve a high degree of crystallinity. The lattice tends to be completely planar, which is in good agreement with the observed flat assemblies from purified CA^CTD^-SP1 proteins (21). To characterize the crystallinity, we computed a two-dimensional (e.g., in-plane) pair distribution function (*g*_dimer_(*r*)) for the dimer center-of-mass, which is shown in Figure 2(b). The observation of distinct peaks is indicative of crystallinity; by contrast, the *g*_dimer_(*r*) in the absence of RNA gradually increases toward unity, which indicates the absence of an ordered structure. Given that the repeating unit in the lattice is comprised of six sub-units of CA-SP1 dimers, the lattice constant can be approximated from the position of the third *g*_dimer_(*r*) peak (refer to Figure S1). Our calculated lattice spacing of 7.4 nm, which is in excellent agreement with previous cryo-EM studies of tubular Gag constructs (~7.25 nm) (18), and reproduction of hexagonal symmetry further support the validity of our CG model.

Visualization of the simulation trajectories (Movie S1) reveals that extended Gag oligomerization proceeds through a multi-stage process. The tethering to RNA seems to increase the local concentration of Gag, thereby facilitating oligomerization. Interestingly, Gag polyproteins continuously undergo cyclical association and dissociation behavior until a hexamer is formed. In turn, these hexamers remain associated indefinitely and serve as nucleating sites for additional hexamer formation, thereby growing the extended lattice. Complementary evidence is provided by the time series profile (i.e., gray line) of the number of Gag (*n*_Gag_) within hexamers shown in Figure 2(c); here, *n*_Gag_ evolves in a step-wise and monotonically increasing fashion. Similarly, fluorescence studies have suggested that Gag recruitment at membrane puncta is irreversible once complete (39). The preferential stability of hexamers points to its importance as a building block of the immature lattice, which we will discuss further in a later section.

While viral RNA seems to orchestrate Gag multimerization, the influence of non-viral RNA on the self-assembly process remains an open question. Non-silencing microRNAs (miRNA) are particularly interesting as a previous study (31) demonstrated their negative regulatory effect on viral infectivity. We proceeded to include short (20-nt) RNA into the CG model to represent non-silencing miRNA and repeated the same simulation procedure as above. Here, we assume no differences between the Gag binding affinities of the viral RNA and miRNA. Hence, the length and concentration of the two RNA species are the only distinctions; the miRNA remains sufficiently long to bind to multiple diffuse Gag (around 1-3) and across Gag hexamers. The time series profiles of *n*_Gag_ for varying concentrations of miRNA are shown in Figure 2(c). Interestingly, the profiles for small concentrations of miRNA (less than a ratio of 200:288:1 miRNA:Gag:RNA) remain consistently above the shaded region, which indicates that the rate of Gag oligomerization is enhanced compared to the miRNA-absent case. However, beyond a critical concentration of miRNA (more than a ratio of 1500:288:1), lattice assembly appears virtually stagnant. Our trajectories show that miRNA-bound Gag impede viral RNA binding, thereby preventing the tethering activity of viral RNA.

Our CG simulation results are supported by evidence from single-particle tracking photoactivated localization microscopy (spt-PALM). We are specifically interested in changes to the diffusive behavior of Gag under different RNA environments since these can manifest due to changes in the multimerized Gag cluster size. We prepared wild-type (WT) and miRNA-overexpressing (has-miR-146a) HEK293 cells transfected with a full-length HIV-1 proviral clone and its derivative tagged with mEOS2, which were expressed in a 3:1 ratio to ensure viral particle formation (see Methods). To remove RNA influence on Gag multimerization (40-42), WT cell lines with NC-deleted Gag (ΔNC) mutants were similarly prepared. Figure 3(a) displays the distribution of Gag diffusivities (*D*) from the WT, miRNA, and ΔNC lines with calculated mean values of 0.092, 0.144, and 0.438 μm^2^/s, respectively. Similarly, a distribution of *D* for multimerized Gag and diffuse Gag dimers as computed from our CG MD simulations are shown in Figure 3(b) with calculated mean values of 570 and 2950 μm^2^/s, respectively. We normalize these two distributions to establish a basis for comparison to account for the well-known separation between CG and real-time dynamics (43, 44); although the CG dynamics are several orders of magnitude faster than reality, the relative ratio of the mean *D* for ΔNC to WT lines (~ 4.8) is comparable to that of CG predictions for oligomerized to diffuse Gag (~ 5.2). It is also notable that the distributions of *D* for the WT and miRNA Gag (Figure 3(a)) and that of oligomerized Gag (Figure 3(b)) exhibit a qualitatively similar preference for slow *D* (< 25% of the mean) while the distributions of both ΔNC and diffuse Gag show comparably broad (i.e., heavy-tail) behavior. These shared features indirectly suggest that Gag can oligomerize in both miRNA-rich and miRNA-absent environments while ΔNC-Gag expectedly remain largely dissociated.

**Figure 3.**
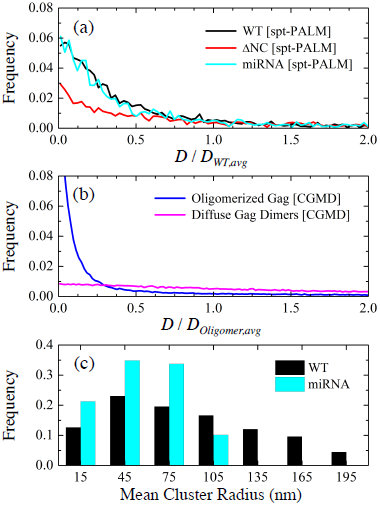
Dynamics and clustering behavior of Gag. (a) Histogram of Gag diffusivities (*D*) from spt-PALM trajectories in WT, ΔNC, and miRNA lines (from 3732, 3619, and 2373 tracks, respectively). These results are normalized by the mean diffusivity of WT Gag (*D*_*WT,avg*_ = 0.092 μm^2^/s). (b) Histogram of *D* from CG MD trajectories in simulations with oligomerized Gag and diffuse (*i.e.*, non-interacting) Gag. These results are normalized by the mean diffusivity of oligomerized Gag (*D*_*Oligomer,avg*_ = 570 μm^2^/s). (c) Histogram of mean cluster radii from spt-PALM trajectories in WT and miRNA lines. Clustered Gag (see Methods) account for 54.2 and 50.9% of tracks, respectively. Bin sizes of 0.0025 μm^2^/s, 10 μm^2^/s, and 30 nm were used in panels (a), (b), and (c), respectively.

We next used PALM cluster analysis (see Methods) to quantify the distribution of Gag cluster sizes in the WT and miRNA lines; ΔNC lines were also analyzed but yielded a significantly smaller fraction (< 10%) of tracks participating in clustering and therefore were not shown. The distributions shown in Figure 3(c) confirm that Gag can form tightly clustered regions in both WT and miRNA lines. However, it is apparent that Gag in WT lines tend to form much larger clusters (approaching 150 nm) compared to miRNA lines. Therefore, our combined simulation and experimental results suggest that while viral RNA instigates Gag multimerization, an excess of miRNA can limit the extent of multimerization and thereby reduce viral particle production (31).

#### Weak Gag-Gag Interaction Strength Regulates Against Kinetic Trapping

To this point, we have established that CA^CTD^-SP1 is assembly competent and nucleates through six-helical bundling at the CA-SP1 junction interface. Importantly, RNA and the plasma membrane also appear to catalyze assembly, which may be a consequence of the weak protein-protein interactions. We next sought to determine if this inherent interaction strength is important for successful assembly by modulating *E*_SP1_ (i.e., the junction binding affinity).First, it is informative to analyze the stability of different Gag oligomers, each observed during general assembly, as *E*_SP1_ is adjusted. To do so, we approximate the probability (*f*_oligomer_) that each of four representative oligomers – a hexamer of dimers (HOD), a closed trimer of dimers (TOD), a chain of TODs, and a crescent fan of TODs (depicted in Figure 4(a)) – resists dissociation due to thermal fluctuations at 310 K within a constant period of time (e.g., 1×10^6^ τ); larger *f*_oligomer_ values imply greater kinetic stability and longer lifetimes. For each type of oligomer, we initialized systems of evenly-spaced 5×5×5 CA-SP1 oligomers separated by 25 nm in all directions withina 125×125×125 nm^3^ simulation domain. Simulations were performed in the canonical ensemble (constant *NVT*) over production runs of 1×10^6^ τ with trajectories saved every 2.5×10^4^ τ for analysis. The reported *f*_oligomer_ is averaged over four independent trajectories for each *E*_SP1_ that is varied between 1.5 and 4.5 kcal/mol.

**Figure 4.**
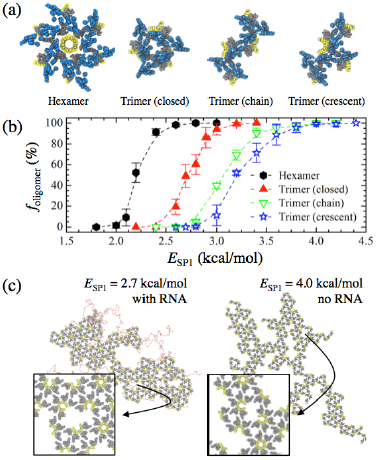
Weak interactions ensure preferential oligomerization into hexamers. (a) Schematic of four representative CA-SP1 oligomers observed during CG MD simulations; blue, gray, and yellow tubes represent helices in the CA^NTD^, CA^CTD^, and SP1 domains, respectively. (b) The probability (*f*_oligomer_) that the listed CA^CTD^-SP1 oligomer remains associated after 1×10^6^ MD timesteps as the SP1-SP1 interaction energy (*E*_SP1_) is varied. (c) Snapshots of final CA^CTD^-SP1 lattices (gray and yellow tubes) assembled at the membrane interface (not shown) when [left] *E*_SP1_ = 2.7 kcal/mol and [right] *E*_SP1_ = 4.0 kcal/mol. In the former (latter) case, RNA (red beads) is still (not) required for lattice assembly to occur. The insets show a zoomed view of the indicted regions, which highlight kinetically-frustrated configurations.

Figure 4(b) shows the *f*_oligomer_ profiles for each oligomer-type as a function of *E*_SP1_. In every case, a sigmoidal profile was obtained in which below (above) a certain *E*SP1, all oligomers spontaneously dissociated (remained associated) while both associated and dissociated states existed in the intermediate *E*_SP1_ regime (i.e., near the inflection point); we should note that the previously used *E*_SP1_ (= 2.35 kcal/mol) is within the intermediate regime for HODs. We find that HODs are considerably more kinetically stable than closed TODs, which in turn are more stable than the open forms of TODs (i.e., chain and crescent forms); note that the tethering of small RNA (20 nt) to each oligomer yielded comparable *f*_oligomer_ profiles. Additional discussion on the factors contributing to oligomer stability can be found in Supporting Information. The preferential stability (i.e., lifetime) of HODs over other oligomers is notable as it further supports the idea of a multi-stage nucleation process in which HODs are assembled through helical bundling at the CA-SP1 junction before the extended lattice is grown; the growth proceeds in a cascading fashion, either from additional HODs nucleating on the edges of the initial HOD or from assimilation of independently formed HODs.

Next, we investigate the impact of increasing *E*_SP1_ to 2.7 and 4.0 kcal/mol, which should increase the stability of closed and open TODs (Figure 4(b)). As depicted in Figure 4(c), the resultant lattice growth exhibits notable anisotropy. This dendritic growth can be attributed to the increased stability of oligomers at the edges of the lattice, which in turn present additional nucleating sites for CA-SP1 consolidation. Furthermore, the degree of anisotropy appears to scale directly with increasing oligomer stability (e.g., the *E*_SP1_ = 4.0 kcal/mol case), while concurrently mitigating the necessity of RNA to promote CA-SP1 association. Vacancy-like defects (i.e., incomplete hexamers) can also form throughout the lattice (see insets of Figure 4(c)). These defects likely arise due to the extended lifetime of non-hexameric oligomers that are subsequently incorporated as kinetically-frustrated configurations. Therefore, unlike our previous simulations (i.e., *E*SP1 = 2.35 kcal/mol), the absence of a self-regulating mechanism to heal lattice defects through cyclic CA-SP1 association and dissociation can result in locally defective immature lattices.

### Immature Lattice Assembly of CA^NTD^-CA^CTD^-SP1 Gag Constructs

#### CA^NTD^ Domain Introduces Natural Curvature through Volumetric Occlusion

In this section, we describe the impact of the CA^NTD^ domain on the structure of the immature CA-SP1 lattice. First, we re-examine the kinetic stability of CA-SP1 oligomers once the explicit volume occupied by the CA^NTD^ domain is introduced. The *f* _oligomer_ profiles depicted in Figure 5(a) exhibit sinusoidal profiles similar to that of the CA^CTD^-SP1 cases (Figure 4(b)). Importantly, the CA^NTD^-CA^CTD^-SP1 profiles are shifted to the right (e.g., the inflection point of each profile is as much as 0.3 kcal/mol higher) in comparison to the previous CA^CTD^-SP1 cases. This indicates that the steric occlusion by the globular CA^NTD^ region tends to destabilize each oligomer. The volume occupied by the CA^NTD^ domain has an important influence on the immature lattice structure. We assembled a hexameric lattice of CA^NTD^-CA^CTD^-SP1 (described inCTD a later section) and computed *g*_dimer_(*r*) profiles between the center of masses of the CA^CTD^ domains and the CA^NTD^ domains for comparison. From Figure 5(b), the sharp peaks associated with the CA^CTD^ and CA^NTD^ layers are indicative of crystallinity. However, one notable difference is their respective lattice spacings, i.e., the position of their third peaks; the CA^CTD^ sub-lattice (~ 7.7 nm) is more tightly packed than the CA^NTD^ sub-lattice (~ 8.5 nm). Given that the lattice spacing of the CA^CTD^ domain is reduced to around 7.4 nm when the CA^NTD^ domain is deleted, these results indicate that the space filled by the CA^NTD^ domain introduces lateral strain to the assembled complex. As the two domains are connected, this strain has the important consequence of introducing an inherent lattice curvature. Following the geometric relationship depicted in Figure 5(c), we approximate the natural radius of curvature (*r*_curv_) to be 39.6 ± 1.3 nm. For comparison, experimental VLPs constructed with ΔMA-Gag can have radii as small as 40-45 nm (45, 46), while WT virions, which are composed of full-length Gag proteins, typically have larger radii that vary between 50 and 100 nm (7, 9, 11).

**Figure 5.**
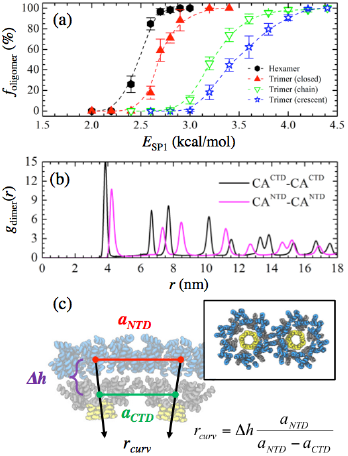
Strain from the CA^NTD^ domain imparts intrinsic curvature. (a) The probability (*f*_oligomer_) that the listed CA^NTD^-CA^CTD^-SP1 oligomers remain associated after 1×10^6^ MD timesteps as the SP1-SP1 interaction energy (*E*_SP1_) is varied. (b) Two-dimensional pair distribution function between the center of masses of the listed Gag domain (g_dimer_) as a function of radial distance (*r*). (c) Schematic of two adjacent CA^NTD^-CA^CTD^-SP1 hexameric bundles depicted as blue, gray, and yellow tubes, respectively. The intrinsic radius of curvature (*r*_curv_) is estimated from the lattice spacing between the center of masses of the CA^NTD^ (*a*_*NTD*_) and CA^CTD^ (*a*_*CTD*_) domains and from the separation distance (?*h*) between domains. Our calculations estimate *r*_curv_ to be 39.6 ± 1.3 nm.

#### Spontaneous Membrane Curvature Triggers Extended Assembly

Surprisingly, we found that CA^NTD^-CA^CTD^-SP1 dimers remained largely dissociated when following our previous simulation procedure (with *E*_SP1_ = 2.6 kcal/mol). This remained true even when *E*_SP1_ was enhanced to 3.5 kcal/mol in an attempt to promote assembly (see Figure S2); although small oligomers readily formed, extended assembly was not observed. We attribute the impeded assembly to the competition between the natural curvature of the CA-SP1 lattice and the resistance to curvature from the membrane. In other words, our simulations suggest that the weak binding affinity between Gag-Gag and membrane-Gag may be insufficient to overcome the free energy penalty for membrane bending. Nonetheless, we found that inducing localized curvature (i.e., small punctas) on the membrane, e.g., from lateral phase segregation of lipids or membrane rafts (47, 48), allowed extended lattice formation to proceed. To further investigate the influence of spontaneous membrane curvature, we instigated local puncta of variable size with a weakly repulsive spherical potential (see Methods) that was slowly pushed (over 5×10^6^ τ) into the membrane with 64 bound dimers and without RNA (with lateral dimensions of 70×70 nm2); we limited system sizes to study a wide range of puncta heights (with respect to the base of the membrane) and radii of curvature. Then, a 2500 nt chain of RNA was introduced and the systems were allowed to equilibrate for 1.5×10^8^ τ at 310 K; it is notable that simulations required much longer trajectories than the CA^CTD^-SP1 cases as the CA^NTD^ domain introduces additional barriers associated with reorganization that slow HOD formation.

Figure 6 summarizes the results of our simulations for CA^NTD^-CA^CTD^-SP1 assembly as a function of puncta height and radius of curvature. Here, Gag self-assembly proceeds at the center of the domain where the membrane puncta (not shown) is located. Interestingly, the efficiency of Gag association, as characterized by the total number of completed hexamers at the end of our simulations, appears to be sensitive to both the height and curvature of the puncta. It is intuitive that the final number of hexamers increases with height as the available surface area of the deformed region also increases. Yet unexpectedly, an optimal radius of curvature also appears at each discrete height, decreasing from around 52.5 to 47.5 nm as the height increases from 2.5 to 7.5 nm. In instances both above and below this critical radius, Gag multimerization appears to be hindered. These results suggest that the alignment between local membrane and intrinsic Gag curvature becomes increasingly important as the lattice grows in order to promote Gag binding onto the cluster periphery. We therefore speculate that the extent of lattice growth is coupled to the dynamics of membrane deformation (i.e., bud formation).To emulate initial budding, we performed a non-equilibrium MD simulation (see Movie S2) of 144 dimers with RNA and a gradually deforming membrane (initial lateral domain size of 100 × 100 nm^2^) over 2 × 10^8^ τ. We monotonically increased the height and radius of curvature of the puncta following the time series profiles shown in Figure 7(a). Figure 7(b) presents *n*_Gag_ profiles according to three classifications: (i) total oligomerized, (ii) within the largest contiguous cluster, and (iii) within complete hexamers. Throughout the trajectory, we observe relatively rapid binding and unbinding of Gag, reined in by RNA, to the central multimerized cluster in order to complete hexamers at the lattice periphery (see Figure 7(c)). Meanwhile, hexamer creation increases monotonically, which is consistent with our observations from TIRF experiments (Figure S3); note that the actual time required for hexamer formation can vary widely as evidenced by the plateaus in the hexamerized *n*_Gag_ profile. The snapshots in Figure 7(c) also show that the lattice grows isotropically such that hexamer formation mostly occurs in concentric layers around the center of the cluster. We can expect this behavior since the entropic burden for hexamer formation (through concerted Gag localization) is reduced for every adjacent hexamer, which provide tethered monomers for hexamer incorporation. Altogether, our simulations suggest that slow and self-correcting Gag assembly concurrent with gradual membrane deformation may be essential features of immature lattice growth.

**Figure 6.**
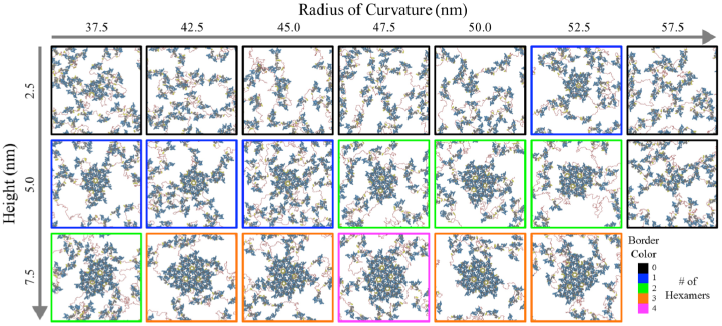
Matrix of final configurations of CA^NTD^-CA^CTD^-SP1 dimers, depicted as blue, gray, and yellow tubes, with RNA (red spheres) at the membrane interface (not shown) after 1.5×10^8^ CG MD timesteps; each snapshot represents an independent simulation. The membrane was spontaneously deformed with a weak spherical indenter (applied centrally) at the listed height and radius of curvature. The border color indicates the number of completed hexamers.

**Figure 7.**
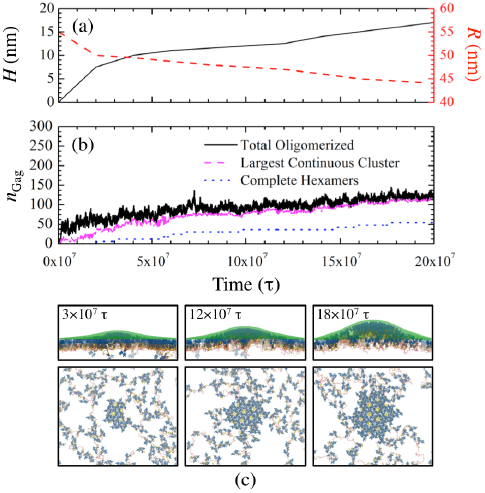
Assembly dynamics of CA-SP1 with RNA at membrane puncta. Time series profiles as a function of MD timestep (τ) of the (a) height (*H*) and radius of curvature (*R*) of the continuously deforming membrane site and (b) the total number of multimerized Gag (*n*_Gag_) categorized in three contexts: anywhere within the simulation domain, within the largest continuous cluster of Gag, and within any completed hexamers. (c) Side and top views of assembled CA^NTD^-CA^CTD^-SP1 lattices, depicted as blue, gray, and yellow tubes, at the listed τ with RNA (red beads) and membrane (transparent green beads) also shown.

## DISCUSSION

We have developed a coarse-grained (CG) computational model from experimental structural data that recapitulates assembly of HIV-1 immature lattices having contiguous and hexameric structures, in general agreement with previous cryo-electron microscopy (cryoEM) and tomography (cryoET) experiments (7-11). Our joint CG molecular dynamics (MD) simulations and single-particle tracking photoactivated localization microscopy (spt-PALM) experiments support recent fluorescence experiments (39, 49-51) that suggested Gag multimerization occurs at the cell membrane interface rather than through the delivery of small Gag oligomers pre-assembled within the cytoplasm. Importantly, we highlight the catalytic role of both membrane puncta and viral RNA to drive multimerization. Our simulations indicate that assembly proceeds through a slow, self-regulated and multi-stage process in which Gag hexamers are gradually formed along the periphery of a growing cluster; the multi-step nature of this assembly process may contribute to the pauses that have been observed during viral assembly (52). We should emphasize that our CG model is intentionally simplified to reconstitute the minimal features required for assembly and may not be directly comparable to the complex ecosystem within cells. As a result, some possible mechanistic nuances, such as molecular conformational switches (21, 22, 24, 30) or regulatory signals from viral RNA-specific sequences and structures (53-56), are not included at this juncture in the modeling. Nonetheless, the simplicity of the present CG model provides a platform to distinguish possible regulatory roles of Gag protein domains, RNA, and the cell membrane on the early stages of immature lattice assembly.

The immature lattice consists of hexagonally-arranged building blocks which themselves are hexamers of Gag. Therefore, successful assembly is contingent on controlled hexagonal order both within the hexameric unit and throughout the extended lattice. Our simulations show that anisotropic attractive interactions at the helical CA-SP1 junction, which may arise from interactions between hydrophobic sidechains (21) that emerge due to the amphipathic character of the helix (22), can provide hexagonal order within the hexameric unit through six-helical bundling. The inherent weakness of this interaction may be significant for two reasons. First, hexamers are preferentially stable over other small oligomers, which subsequently allows non-ideal oligomers to anneal into proper hexamers. Secondly, hexamer formation requires concerted localization and reorganization of Gag, which can be facilitated by the RNA scaffold that, in part, makes RNA indispensable (and ensures that RNA is packaged). Based on our simulation results, we also hypothesize that enhancing the interactions at this helical interface can either induce aberrant Gag multimerization or abolish RNA packaging requirements; for example, A366E and M367K mutations exhibit Gag localization on the plasma membrane (i.e., arrested virion release) or non-spherical virion phenotypes (22), which may be interpreted based on additional electrostatic attraction at the helical junction.

Once the formation of the initial hexamer unit is complete, additional hexamers nucleate in concentric rings around this seed. Our results suggest that this process is enabled by both Gag dimerization and Gag recruitment driven by RNA. Recall that Gag dimerization, which occurs through the H9-H9 interface in the CA^CTD^ domain, is preferred in the cytoplasm (57), although the relative orientation of these helices differs in mature CA dimers and lattices (18, 58-60). Hence, hexamer formation utilizes six distinct dimers but only half of each; the other half of each dimer remains flexibly bound on each of six facets. These dangling dimers serve as a nucleating template by reducing the number of Gag that must be recruited through RNA scaffolding; as a result, the immature lattice can expand in a largely contiguous fashion (8, 9). Based on our findings, we suggest that efficient particle release may require RNA that is sufficiently long to continuously recruit free Gag into the lattice. Previous experiments have demonstrated that viral RNA as short as 3 kb can generate viral particles (61), although the efficacy of much shorter RNA remains to be explored. Our joint experiments and simulations on ultra-short (20-nt) miRNA, and related observations from previous experiments using miRNA (31) and tRNA (30), suggest that an excess of short competitor RNA can negatively regulate the ability of viral RNA to orchestrate Gag multimerization by restricting the availability of Gag-RNA binding sites.

The dynamics of membrane deformation during budding appear to further regulate the assembly process. This is related to our finding that the CA^NTD^ domain imparts intrinsic curvature to the immature lattice. There has been some debate on the role and relevance of the CA^NTD^ domain since ΔCA^NTD^ Gag has been reported to successfully produce VLPs (27, 46), remain aggregated at the plasma membrane (62), or assemble into flat sheets *in vitro* (21). Our findings show that both CA^NTD^-Gag and ΔCA^NTD^-Gag are assembly competent, but the former may require spontaneous membrane curvature to initiate nucleation while extended growth is contingent on gradual membrane deformation. Beyond simple thermal fluctuations, curvature may be passively promoted by the heterogeneous compositions of membranes, such as through lipid raft domains (63-65) or transmembrane proteins (66-68). In fact, the binding of MA domains (and associated myristyl anchors) to the membrane may be important to initiate PIP_2_ enrichment into nanodomains (36, 69) and possibly membrane bending. In addition, the actin cytoskeleton has been proposed to expedite viral assembly and budding (70), although its role is dispensable as shown by recent fluorescence experiments (71, 72). We further speculate that membrane deformation dynamics can influence the formation of previously observed defects throughout the immature lattice (8-10), especially since immature virions tend to be highly pleomorphic. While local hexameric defects may be similar to the vacancy-like defects observed in this work, extended gap-like defects might also arise from anisotropic lattice growth at the leading edges, which might be instigated by the strain imposed from the mismatch between the local curvature of the Gag lattice and membrane. Hence, the presence of these possible curvature generation factors is likely important to understand differences in observed virion phenotypes.

To summarize, our findings reveal that RNA and the cell membrane are active participants of viral assembly that both initiate and coordinate Gag multimerization. Deformations along the membrane can regulate the location and size of the assembling Gag cluster, which can only proceed when RNA drives the localization and reorganization of multiple free Gag toward the cluster periphery. Our results suggest that immature lattice assembly and budding are coupled and proceed concurrently (73). These insights reveal a network of coordinated interactions that are likely to be essential for viral replication.

## METHODS

### CA-SP1 Coarse-Grained Model

To reproduce the internal structure of the CA-SP1 dimer, we only considered the structured portions within each protein domain. As depicted in Figure 1(a), the relative position of each CG site was obtained from the average position of each C-α throughout the α-helices within the CA^NTD^ (7 helices), CA^CTD^ (4 helices) and SP1 (1 helix) domains, which were calculated over all monomer chain configurations. We used two experimental data sets (PDB: 4USN and 5I4T) that were superposed by minimizing the root-mean-squared deviation of the C-α positions in the CA^CTD^ domain. As NMR spectroscopy has suggested that each protein domain remains folded in solution and reorients semi-independently from each other (57), the collection of α-helices within each domain was considered as a rigid body with rigid body dynamics (74). The tertiary structure of CA-SP1 was maintained with an elastic network model (ENM) that connected CG particles in adjacent domains (e.g., between helix 10 of CA^CTD^ and the SP1 helix) through flexible harmonic bonds; the potential energy of the bonds (*E*_bond_) was given by

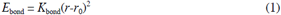

where *K*_bond_= 0.5 kcal/mol/Å^2^, *r* is the separation distance, and *r*_*0*_ is the computed average distance. To maintain the shape of the polyprotein, each CG particle had an excluded volume through a soft cosine potential (*E*_excl_) given by

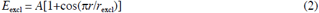

when *r* < *r*_excl_. The excluded volume separation (*r*_excl_) between CG particles in adjacent helices was set to their average separation distances if less than 10 Å or otherwise, set to 10 Å (i.e., CG particle radii are 5 Å by default); in all cases, *A* = 25 kcal/mol. The mass of every CG site was set to 150 Da.

The CG model also incorporated two important protein-protein interfaces that have previously been identified. First, we used a flexible ENM (*K* _bond_ = 0.05 kcal/mol/Å^2^) between adjacent helix 9 (H9) in the CA^CTD^ domains to dimerize CA-SP1 based on the helical orientations observed in the aforementioned experimental data sets; our ENM represents the strong preference for CA proteins to exist as homodimers in the cytoplasm (57, 60), although the relative tilt between H9 can vary (58, 59). In addition, we assume that the CA-SP1 junction forms an ordered helix that assembles into a high-density bundle (8, 10, 18-21). To recapitulate this interface, we used the relative positions of four residues (P356, A360, A364, S368) in an adjacent SP1 domain when confined within a hexamer configuration to assign the positions of virtual particles (pink spheres in Figure 1(a)). Each of these virtual particles were allowed to tether to any CG particle of the same residue through an attractive Gaussian potential (*E*_tether_) given by

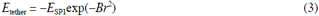

where *E*_SP1_ is the well-depth (varied throughout this manuscript), *B* is the inverse interaction range (= 0.2 Å^−2^), and *r* is the distance between the virtual and tether site. This interaction was the only attractive non-bonding interaction between CA-SP1 dimers used throughout this work and effectively represents the net contributions from the SP1 helical interactions, the major homology region, and the nearby type II β turn (19-21).

By combining the excluded volume and structural ENM throughout the CA-SP1 polyprotein, the flexible ENM at the H9-H9 dimer interface, and the anisotropic tethering potential at the CA-SP1 junction, the tertiary and quaternary structure of assembled CA-SP1 recapitulated the expected hexamer structure (see Figure 1(b)).

### RNA and Membrane Model

Simplified CG models of RNA and the membrane were used throughout this work. RNA was represented as a linear chain of CG particles that were linked with flexible harmonic bonds (*K*_*bond*_ = 0.5 kcal/mol/Å^2^) and excluded volume with *r* = 7.0 Å (i.e., particle radius of 3.5 Å). The mass of each CG site was set to 330 Da. To implicitly include binding between RNA and the NC domain, each RNA particle was allowed to tether to the final SP1 particle with a well-depth of 7.5 kcal/mol and inverse interaction range of 0.1 Å^−2^; as this interaction is, in part, due to electrostatic attraction between the negatively charged phosphate groups in RNA and positively charged Zn^2+^ fingers in the NC domain (53, 75), the chosen potential energy is intentionally both stronger and wider-ranged than that of the hydrophobic SP1-SP1 interaction (21, 22).

The simplified membrane was represented by an elastic mesh of particles (with a mass of 1 kDa) arranged in a hexagonal lattice with a lattice constant of 0.85 nm. Bonds between particles are maintained with a Morse potential (*E*_morse_) given by

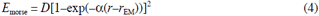

where *D* = 450.0 kcal/mol, α = 0.17 Å^−1^, and *r* = 5.0 Å; each particle was connected to their three neighbors. Particles also interacted through a long-range 12-6 Lennard Jones potential with a modified soft core (*E*_sclj_) (76) given by

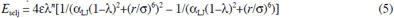

where *n* = 2, α_LJ_ = 0.5, λ = 0.6, ε = 0.03 kcal/mol, and ρ = 20 Å. These parameters have been adjusted to reproduce a fluctuation spectrum (see Figure S4) with bending modulus of around 12 *k*_*B*_*T* (77). Our intention was simply to emulate the membrane as a substrate that reduces the dimensionality of CA-SP1 assembly (65). To implicitly include binding between the membrane and Gag through exposed myristyl and the MA domain (78, 79), each elastic mesh particle interacted with the top-most particle in helix 4 of the CA^NTD^ domain through *E* _sclj_ where *n* = 2, α_LJ_= 0.5, λ = 0.6, ε = 1.5 kcal/mol, and ρ = 12 Å; note that weaker interaction strengths prevent Gag binding while stronger interaction strengths prevent extended lattice growth (see Figure S2) In addition, spontaneous curvature in the membrane was induced through a spherical indentation force (*F*_ind_) given by

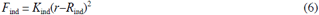

where *K*_ind_= 0.001 kcal/mol/Å^3^ and *R*_ind_ is the radius of the indenter; the force is repulsive when *r* < *R*_ind_ and zero otherwise. In these cases, an opposing planar force (*K*_ind_= 0.0001 kcal/mol/Å^3^) was weakly applied to the outermost edges of the mesh (with thickness around 15 Å) to prevent vertical drift.

Unless otherwise specified, default excluded volume interactions (e.g., radius of 5 Å) were maintained between CA-SP1, RNA and the membrane.

### Molecular Dynamics Simulations

All CG MD simulations were performed using the Large-scale Atomic/Molecular Massively Parallel Simulator (LAMMPS) (80). Three different classes of simulations were performed throughout this manuscript; the general details of each are described below. In all cases, a temperature of 310 K was maintained using Langevin dynamics (81) with a damping constant of 100 ps. A Nosé-Hoover chain barostat (82, 83) in the coupled *xy* (i.e., lateral) dimensions was used to maintain zero tension with a damping constant of 250 ps, unless otherwise specified. A CG MD timestep (τ) of 200 fs was used; we chose the largest τ that was possible without noticeable energy drift in the microcanonical (constant *NVE*) ensemble. All pair potentials used a 25 Å radial cutoff with an additional 5 Å buffer for particle neighbor lists. All simulations were periodic in all three directions although in all cases, unless otherwise specified, the *z* direction was bound by repulsive walls and essentially non-periodic. Several different simulations were performed throughout this manuscript; specific details are described within the appropriate subsections in Results.

### Oligomerization Metrics

Gag oligomer sizes were determined through a recursive search for minimum path lengths over a directed graph. Cyclical paths with length 6 were counted as hexamers while non-cyclic paths with length less than 6 (while greater than 0) were counted as smaller oligomers. To construct the directed graph, an edge was formed between two SP1 helices (from different dimers) whenever all four SP1 virtual sites of one helix were within 0.2 nm of their respective binding sites on the other helix.

### Pair Distribution Functions

Two-dimensional (i.e., in the *xy* plane) pair distribution functions were computed over the center of mass of each Gag dimer; the center of mass of the CA^NTD^ and CA^CTD^ domains were used for further demarcation. Statistics were gathered for 1×10^7^ τ with data printed every 5000 τ. At each frame, the number of Gag within a *xy*-distance of 20 nm (using 0.1 Å bins) was computed and distributed into 0.1 Å bins with normalization to the area of the radial shell. The reported pair distribution functions were obtained from the ensemble averaged distributions normalized by the average density of Gag within 20 nm.

### Diffusivities

Mean squared displacements (MSD) were computed based on the center of mass of each Gag dimer. Statistics were gathered for 1.5×10^7^ τ with data printed every 5000 τ. Particle trajectories over 15 frames within overlapping time windows were used to compute MSD as a function of CG time; computed MSDs were two-dimensional as all positions were projected onto the *xy* plane. Linear regression was performed to compute the slope of the MSD profiles (*S*_MSD_) for each trajectory. The diffusivity (*D*) from each trajectory was then computed from the Einstein relation (*D* = *S*_MSD_/4).

### Preparation of Cells and Plasmids

Wild type HEK293 cells and HEK293 cells overexpressing exogenous human microRNA (miRNA) has-miR-146a were cultured and maintained in phenol red free Dulbecco’s modified eagle medium (DMEM) supplemented with 10% (vol/vol) FBS (Invitrogen) and 2mM glutamine (Cellgro, Herndon, VA). Designs of miRNA expression plasmids and viral constructs used for expressing Gag (pNL4-3ΔPΔE, pNL4-3-imEOS2-ΔPΔE and pNL4-3-iEGFP-ΔPΔE) and generation of miRNA cell line have been described before (31).

### PALM Imaging and Analysis

Live cell PALM experiments were carried out with an Elyra PS.1 using a 100× 1.46 NA oil immersion objective (Carl Zeiss, Thornwood, NY). Cells were plated in cleaned 25mm #1.5 coverslips (Warner Instruments) and transfected with pNL4-3ΔPΔE and pNL4-3-imEOS2-ΔPΔE plasmids as described before (31). Live cell single particle tracking PALM (spt-PALM) experiments (31, 37) were performed with cells placed in imaging medium (phenol red free DMEM containing 25 mM Hepes and 1% FBS) maintained at 37°C. During PALM image acquisition, mEOS2 probes were photoconverted using a 561 nm laser (readout induced activation mode) and images were collected at 20 frames per second. The 561 nm laser power was calibrated to maintain a sparse distribution of single mEOS2 fluorescent spots at every image frame of the time series.

PALM images of live cells were collected using illumination conditions where the average distance between single molecule fluorescent spots was significantly larger than the localization precision of single Gag molecules. Single mEOS2-tagged Gag molecules were identified and fit with a two-dimensional symmetric Gaussian point spread function using either commercially available software (Zeiss Zen Black super resolution module, Carl Zeiss) or custom code written in MATLAB. Only single molecule peaks with localization precision less than 35 nm were used for analysis; the average localization precision of the peaks ranged between 20 and 35 nm. The maximum distance that a single molecule can traverse between two consecutive frames depends on the average diffusion coefficient of the molecule. Previous studies have reported that the maximum diffusion coefficient of Gag molecules on mammalian plasma membrane is ~ 0.1 μm^2^/s. This implies that 99% of Gag-molecules would move less than 300 nm in the time period (50 millisecond) between two consecutive image frames. Using this information, we assigned single molecule peaks in consecutive frames to the same trajectory if the distance between the peaks was less than 300 nm. The significantly large average distance between single molecules within an image frame (around 3 μm) made it highly unlikely that two peaks in consecutive frames within 300 nm of each other represent positions of distinct molecules. This algorithm was used to stitch single molecule positions across the entire PALM image dataset and generate trajectories of single molecule diffusion. Each trajectory was assigned a unique track ID. Next, trajectories containing at least 12 steps were identified. For each trajectory, the average mean square displacement (MSD) of the molecule for different time lags (Δt) was calculated by averaging the MSD across overlapping time windows. Finally, linear fitting of the MSD vs. Δt across the first five time lags (0 < Δt < 250ms) was used to calculate the average short-term diffusion coefficient for each trajectory.

The spatial organization of the localized single molecules was analyzed by performing Hoshen-Kopelman algorithm-based cluster analysis (84) of the live cell PALM data set. First, a composite super-resolution image was generated by combining all the single molecule positions identified in the entire set of image frames of a PALM time-series experiment. The composite image was analyzed using the Hoshen-Kopelman algorithm to identify and spatially localize individual clusters of Gag molecules. All neighboring Gag molecules within a distance of 150 nm of each other were recursively assigned to the same cluster. Next, the average density of Gag molecules in each cluster was calculated. Only clusters with density greater than three times the average density of Gag over the plasma membrane were considered as Gag platforms representing nascent viral buds and were assigned a unique cluster ID. Next, in order to estimate the dimensions of each Gag cluster, the convex hull for the set of molecules comprising the Gag cluster was calculated. The area of the convex hull was used as an estimate of the area of the Gag cluster. The radius of a circle with the same area as the convex hull gave an estimate of the cluster radius.

## ACKNOWLEDGEMENTS

This work was supported in part by the National Institute of General Medical Sciences of the National Institutes of Health under award number P50-GM082545 (G.A.V, M.Y.). The Lippincott-Schwartz laboratory acknowledges financial support from the Howard Hughes Medical Institute (HHMI). The Briggs laboratory acknowledges financial support from the European Molecular Biology Laboratory (EMBL), Chica und Heinz Schaller Stiftung, and Deutsche Forschungsgemeinschaft grants BR 3536/2-1 (J.A.G.B.). Computational resources were provided by the Blue Waters sustained-petascale computing project, which is supported by the National Science Foundation (awards OCI-0725070 and ACI-1238993) and the state of Illinois. Blue Waters is a joint effort of the University of Illinois at Urbana-Champaign and its National Center for Supercomputing Applications. This work is part of the ‘Ultra-Coarse-Grained Simulations of Biomolecular Processes at the Petascale’ Petascale Computing Resource Allocation (PRAC) support to G.A.V. by the National Science Foundation (award number OCI-1440027).

## Supporting Information

### Movie S1

Assembly of CA^CTD^-SP1 Gag dimers into an extended lattice through RNA tethering at the membrane interface.

### Movie S2

Assembly of CA^NTD^-CA^CTD^-SP1 Gag dimers into an extended lattice at a membrane puncta through RNA tethering and gradual membrane deformation.

**Figure S1.**
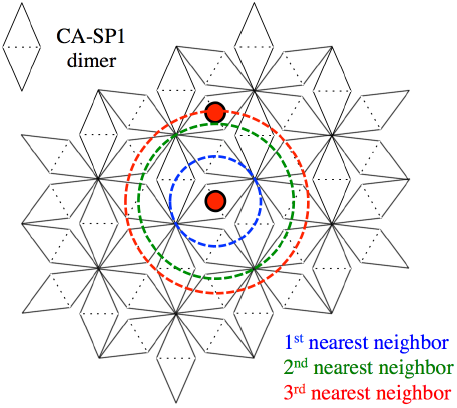
Schematic representation of the assembled immature lattice from CA-SP1 dimer sub-units. The dashed lines indicate the dimerization interface; hexagons bounded by the dashed lines are hexamers. The two red circles indicate the center of dimers in adjacent hexamers; the distance between the two red circles is the lattice spacing. The three dashed rings show the first three nearest neighbor shells. It can be seen that the lattice spacing is equivalent to the 3^rd^ nearest neighbor distance (with respect to the center of dimers).

**Figure S2.**
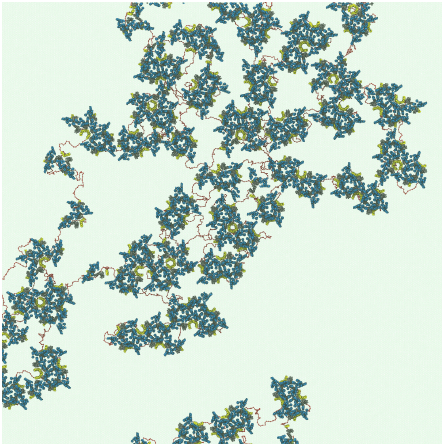
Final snapshot of CA^NTD^-CA^CTD^-SP1 assembly with *E* _SP1_ enhanced to 3.5 kcal/mol after 1×10^8^ MD timesteps; each of the CA^NTD^-CA^CTD^-SP1 domains are depicted as blue, gray, and yellow tubes, respectively, while RNA and the membrane are shown as a red chain and transparent green mesh, respectively. Note that despite the enhanced CA-SP1 interactions, only small oligomers formed while extended lattice assembly was not observed.

**Figure S3.**
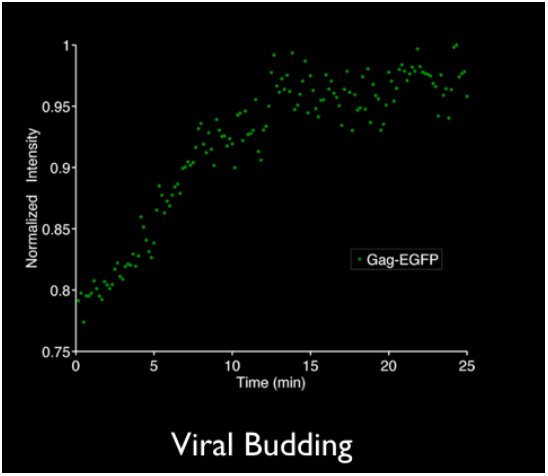
Time series profile of normalized EGFP-fluorescence intensity within a diffraction limited spot during the assembly of a single viral particle. The exponential increase in signal intensity (< 12 min) indicates progressive accumulation of EGFP-labeled Gag to the assembling platform. The plateau in the signal (> 12 min) marks the end of viral assembly.

**Figure S4.**
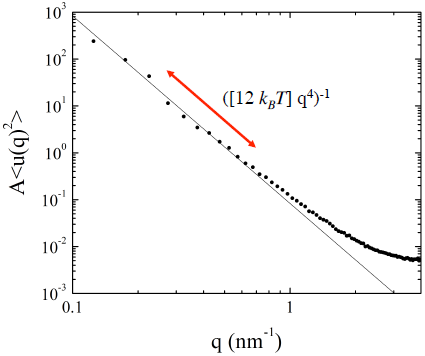
Areal undulation spectrum (A<u(q)^2^>) of the elastic mesh as a function of wave number (q) shown as black dots. The solid line (∝ q^−4^) represents the fit on small-q data used to extract the bending modulus, which is approximately 12 *k*_*b*_*T*. All statistics were gathered from an *NP*_*xy*_*T* simulation using a Langevin thermostat and Nosé-Hoover barostat [see main text] with an integration timestep (τ) of 50 fs. A 100×100 nm^2^ mesh with 29640 CG particles was simulated for 2×10^6^ τ with statistics gathered every 1000 τ.

### Sample Preparation and TIRF Imaging

HeLa cells plated on fibronectin coated glass bottom dishes (MatTek, Ashland, MA) were transiently transfected with plasmids pCR3.1/HIV-Gag and pCR3.1/HIV-Gag-EGFP at a 10:1 molar ratio. TIRF imaging of live cells were performed 6-8 hours post transfection with a customized inverted Nikon-TiE microscope using a 100X Apo TIRF 1.49 NA oil immersion objective (Nikon Instruments, NY) at a frame rate of 10 seconds/frame. Custom MATLAB code and ImageJ macros were used to identify sites of single viral bud assembly (represented by isolated point spread functions on the membrane surface) and quantify the evolution of fluorescence intensity within assembly sites.

### Brief Discussion on Kinetic Stability of Oligomers

Here, we briefly discuss the factors that contribute to the kinetic stability of the Gag oligomers explored in the main text. Let us imagine that each of these oligomers represents a metastable state in free energy space that is separated from the completely dissociated state by an activation barrier of height *E*_a_. One factor that can modulate *E*_a_ is the relative energetics (i.e., internal energy) of the oligomer configurations if we assume the free energies of the transition states are comparable. Given that the oligomers associate through a clear attractive interaction at the SP1-SP1 interface (in our model), we can approximate the energetics of each oligomer through the number of SP1 contacts normalized to the number of Gag dimers; for now, we assume differences in strain energy from excluded volume interactions are negligible compared to this attractive interaction. It is clear that closed oligomers (e.g., HODs and closed TODs) contain a larger number of binding interfaces (*n* SP1 helical contacts per *n* dimers) than open oligomers (e.g., chain and crescent TODs with *n*-1 SP1 helical contacts per *n* dimers). In turn, the more favorable energetics in the former cases can lower the free energy and increase *E*_a_. This explains, in part, the greater kinetic stability observed for closed oligomers compared to open oligomers.

Now despite the similar binding energies between HODs and closed TODs, the former still tends to be more kinetically stable. We reason that the enhanced stability of HODs compared to TODs arises from the bi-directionality of the anisotropic SP1 interaction in the former compared to the uni-directional interactions in the latter. Within HODs, the SP1 domains that participate in binding interact through two interfaces (i.e., two adjacent SP1 helices). In closed TODs, the SP1 domains that participate in binding interact through one interface. It is possible that the bi-directional interactions in HODs restrict the configurational degrees of freedom more so than the uni-directional interactions in TODs. However, at this point, it is unclear if these considerations can increase the free energy of TODs relative to HODs due to entropic differences, increase the free energy of the transition state for HOD dissociation relative to TOD dissociation, or both. Since the kinetics of oligomer association and dissociation are not the focus of this work, we end the discussion here. Nonetheless, unraveling the thermodynamics and kinetics of this self-assembly process could warrant future investigation.

